# Identification of genetic variation associated with high-temperature tolerance in cowpea

**DOI:** 10.1101/2025.04.09.647789

**Authors:** Akakpo Roland, Lee Elaine J, Pacheco Jacob B, Rios Esteban F, Kantar Michael B, Boukar Ousmane, Volz Kevin M, Akinmade Habib, Getino Luis, Boote Kenneth J, Muñoz-Amatriaín María, Morrell Peter L

## Abstract

Heat tolerance is an important trait in cowpea, a crop that constitutes the primary protein source for a large portion of the human population in sub-Saharan Africa. Cowpea grows across this region, with cultivated, landrace, semi-wild, and wild cowpeas germplasm growing across diverse climatic conditions. This study used environmental association (envGWAS) and allele frequency outlier approaches in a panel of 580 gene bank accessions to identify genomic regions associated with heat and limited precipitation. Because allele frequency outliers are detected independent of potential selection factors driving differentiation, we used a ranking-based approach to identify the climate variables most associated with variants among outliers. Precipitation-related variables dominated the signals we identified for envGWAS and allele frequency outliers. We found variants on all eleven chromosomes putatively associated with the adaptation of cowpea to higher-temperature environments. The considerable overlap between variants associated with low precipitation and high temperature suggests that these traits may be inextricably linked in cowpea. The Sahel region is the source of many accessions with derived variants associated with high temperature, suggesting the potential for accessions from this region to contribute to heat tolerance alleles for cowpea improvement.

## Introduction

Ongoing environmental change is affecting crop productivity and the stability of food systems (Wheeler and von Braun, 2013). Rising global temperatures are causing drought and heat to become predominant plant stresses (Pereira, 2016). As a result, strategies are urgently needed to improve plant adaptation to new climate scenarios (Raza et al., 2019). Crop production is vulnerable to changes in climate, with the increase in temperature being among the factors most negatively impacting crop yield. Heat stress occurs when temperatures are hot enough for a sufficient period that they cause either irreversible damage to plant development and function or an increased rate of reproductive maturation, both of which result in a reduction of grain yield (Basra, 2023).

Abiotic stresses influence species distribution across diverse environments (Eckhart et al., 2011; Paes de Melo et al., 2022). Plant adaptation to various environmental conditions in response to abiotic stresses often involves multiple survival strategies, including changes in morphology, physiological adjustments, and molecular reactions (Shafi et al., 2022). These adaptive responses encompass a spectrum of reactions ranging from changes in gene expression and physiological adjustments to plant architecture and modifications in primary and secondary metabolism (Mareri et al., 2022). Efforts to identify the genetic basis of plant adaptation to abiotic stress can provide valuable knowledge about how plants respond to stress, which is crucial for breeding resilient varieties.

An early attempt to identify adaptive differences among populations focused on allele frequency differentiation (Lewontin and Krakauer, 1973). Efforts to identify the genetic basis of agronomically adaptive traits have focused for many years on bi-parental crosses and identifying quantitative trait loci (QTL), contributing to the phenotypic variation in segregating populations (Lander and Botstein, 1989). Advances in association or LD mapping (Lander and Schork, 1994) approaches have bridged these somewhat divergent approaches (Ross-Ibarra et al., 2007). Genome-environment association or environmental genome-wide association studies (envGWAS) have been developed to use environmental summaries to identify variants contributing to adaptation to particular environmental conditions (Bragg et al., 2015; Rellstab et al., 2015; De La Torre et al., 2019). These approaches implicitly assume that measurable environmental variables can be used as proxies for the selective forces shaping genetic differentiation among populations and individuals across geographic space (Tiffin and Ross-Ibarra, 2014). Challenges for applying environmental association include appropriately accounting for population structure (Günther and Coop, 2013) and using multiple environmental measurements that result in highly correlated variables (Hoban et al., 2016). There is also the challenge of multiple environmental factors creating interacting selective pressures that shape allele frequencies in ways that do not accord with individual environmental variables (Lotterhos, 2023). Nonetheless, in barley (*Hordeum vulgare* L.), where many large-effect loci controlling adaptive traits have been identified through QTL and association mapping, envGWAS, and allele frequency differentiation approaches detected most of the known loci (Lei et al., 2019).

Environmental association approaches, combined with complementary analyses such as allele frequency comparisons with F-statistics and comparisons of allele frequency gradients, have been used in several systems to identify genes likely to explain adaptive differences among populations (Pyhäjärvi et al., 2013; Anderson et al., 2016; Neyhart et al., 2022). In common bean (*Phaseolus vulgaris* L.), López-Hernández et al. (2019) examined the genetic control of heat tolerance. Although this study utilized a limited number of wild common bean accessions (78), authors identified candidate genes related to heat stress response, including heat shock proteins. Moreover, the study identified genes involved in multiple biological processes potentially correlated with plant adaptation to high temperatures, including time to flowering, germination and seedling development, cell wall stability, and abiotic stress signaling pathways (López-Hernández et al., 2019). In barley, envGWAS, using bioclimatic variables, identified six well-known loci associated with adaptive variation in flowering time and abiotic stress tolerance (Lei et al., 2019). In maize (*Zea mays* L.), an envGWAS study addressed gene expression responses to various environments across Mexico (Gates et al., 2019). This study identified loci associated with phenotypic and fitness differences across environments, providing insights into local adaptation. In addition to identifying allele frequency differentiation between populations, steep allele frequency gradients have been used to detect adaptive variants (Lasky et al., 2023). Spatial ancestry analysis (SPA) combines locality information and the genotypic state for each sample to identify sharp changes in allele frequency across the landscape (Yang et al., 2012). The direction of allele frequency gradients can differ by variant. This avoids the definition of discrete populations, a long-standing challenge with F-statistics comparisons (Yang et al., 2012).

Cowpea (*Vigna unguiculata* L. Walp.) is an African warm-season legume that provides a valuable source of protein in the diets of millions of people in sub-Saharan Africa and a nutritious fodder for livestock. Cowpea is also one of the crops listed in the International Plant Genetic Resources Treaty for Food and Agriculture (IPGRTFA) Annex I, which falls under the global multilateral benefit-sharing agreements. The species is well-adapted to hot and dry environments. Thus, most of the worldwide production is concentrated in drought- and heat-prone zones of sub-Saharan Africa (Omomowo and Babalola, 2021; Abebe and Alemayehu, 2022; Affrifah et al., 2022). However, relatively high nighttime temperatures can damage reproductive processes, including the flowering pod and seed set, negatively impacting grain yield (Porch and Hall, 2013). Previous studies using commercial cowpea varieties have shown linear grain yield reductions of ∼5% per every 1°C increase in minimum nighttime temperatures above a threshold of 15°C, up to a 50% grain yield reduction at minimum night temperatures of 26°C (Nielsen and Hall, 1985a; Nielsen and Hall, 1985b; Hall, 2004). In addition, Nielsen and Hall (1985a) showed that increases in minimum nighttime temperature from 16°C to 26°C caused a 7-day reduction in the pod development period, which shortens the time to capture resources and reduces yield. These studies were conducted under long-day conditions in the subtropics, where nights tend to be colder than those of the major tropical production zones. The photoperiod-sensitive germplasm, which does not flower under long days, has not been broadly surveyed and could be a reservoir of beneficial alleles for heat stress tolerance.

Extensive genetic resources for cowpea, including landraces and wild relatives, are available from germplasm collections worldwide. These collections harbor functional allelic variation for adaptation to abiotic and biotic stresses. One of the largest cowpea collections is at the International Institute of Tropical Agriculture (IITA), which also holds a Core set of >2,000 accessions that have been genotyped with 51,128 SNPs on the Illumina iSelect Consortium Array (Close, 2023).

GWAS approaches using germplasm collections have identified several loci and candidate genes associated with agromorphological and adaptive traits, including pod shattering (Lo et al., 2021), flowering time (Muñoz-Amatriaín et al., 2021; Paudel et al., 2021), and other phenological traits (Andrade et al., 2023). In addition, QTL mapping in two bi-parental populations reported QTLs underlying heat tolerance on most cowpea chromosomes (Lucas et al., 2013).

In this study, we addressed five questions. First, can environmental association approaches identify variants where allele frequency differentiation suggests adaptation to high temperature tolerance? Second, to what extent are variants associated with high-temperature tolerance independent of those identified for adaptation to water deficit? Third, how often do variants identified using mixed model association methods overlap with those found using *F*_ST_ (allele frequency differentiation) and SPA (allele frequency gradient) approaches? Fourth, to what degree are variants identified in *F*_ST_ and SPA analysis associated with extreme temperature and rainfall values based on environmental descriptors of germplasm collection locations? And fifth, given the patterns of linkage disequilibrium in this cowpea collection, to what degree can we identify potential causative loci from the individual variants surveyed?

## Materials and Methods

### Germplasm and passport data curation

This study took advantage of the IITA Core collection, which includes 2,082 worldwide cowpea accessions that have been genotyped at a high density (Fiscus et al., 2024). We initially focused on 683 landraces and wild cowpea accessions from Africa with locality information, excluding breeding lines and landraces collected at a “market.” Passport data was reviewed in detail, using both manual and automated strategies with the tool ITALLIC (Onsongo et al., 2022). Inconsistencies were found for 190 accessions, mainly related to mismatches between geographic coordinates and detailed location information. In a coordinated effort with IITA gene bank curators, passport data was updated for 59 accessions, while 131 accessions with insufficient data remained unchanged (Table S1). A large set of the latter (98 accessions) seemed problematic, as they were categorized as landraces, but all available information suggested they were breeding lines developed by an IITA cowpea breeder in the 1980’s. These lines were not included in this study. In addition, five other samples were eliminated, mostly because their location information pointed to a research institute. Two additional accessions from Somalia, which previously lacked latitude and longitude information and were therefore excluded from the initial set of 683 accessions, were included in the study as the information was retrieved from their passport data. The curation process resulted in a final count of 580 accessions.

### Verifying geographic origins

Given the curation required to verify the geographic origin of accessions, we sought additional confirmation of origin through the comparisons of genotype-based prediction of the location of origin relative to passport coordinates. We used a deep-neural network tool for spatial ancestry inference (Battey et al., 2020) implemented in the software locator. The inputs for this analysis were iSelect SNP genotyping data and the geographic coordinates for accessions. A randomly chosen set of 16 accessions was treated as a query sample, and the balance of accessions was used for training. Uncertainty of sample placement and a 95% confidence interval for sample origin were estimated using 100 bootstrap replicates per query sample.

### Extraction of bioclimatic data and SNP curation

A total of 580 cowpea landraces and wild accessions from Africa for which GPS coordinates were accurately determined were used. Those GPS coordinates were used to extract historical climatic data available at WorldClim v2 (http://www.worldclim.com/version2) (Hijmans et al., 2005; Fick and Hijmans, 2017). The accessions list originated from 30 African countries, with 1 to 173 accessions per country. Their passport data, including geographic coordinates, are provided in Table S2.

Genotypic data from 51,128 SNPs were available for these accessions and were curated in Plink 1.9 (Chang et al., 2015) to remove SNPs with >20% of missing data and a MAF < 0.02. The physical locations of SNPs were determined relative to the IT97K-499-35 genome assembly (Lonardi et al., 2019) using the approach described in (Liang et al., 2024). SNP locations are available in VCF format at https://github.com/MorrellLAB/cowpea_annotation/blob/main/Results/IT97K-499-35_v1.0/iSelect_cowpea.vcf.

From “WorldClim 2.1”, we extracted bioclimatic data at a resolution of 30 seconds on February 18th, 2023. The getData function from the R package ‘raster’ (Hijmans, 2024) was used to extract the data for BIO1 to BIO19 (Table S3). These 19 bioclimatic variables are potentially redundant. To determine the degree of redundancy, we calculated the correlation between them (Fig. S1). We then summarized the 19 bioclimatic information by performing independent component (IC) analysis using the ‘ica’ R package (Durieux et al., 2024), resulting in three additional variables as considering the three first ICs (Table S4). The independent components (ICs) approach is conceptually similar to principal components analysis in summarizing data. We also used two other temperature variables corresponding to the maximum and the minimum temperatures associated with the approximate flowering period of the crop in each location, as well as the mean precipitation over the same period. These data were extracted from the WorldClim V2 database using the getData function from the R package ‘raster’ (Hijmans, 2024). These three additional variables were obtained by averaging their values over the target flowering period. Altogether, we used 25 environmental variables to identify variants associated with heat tolerance.

### Structure of genetic diversity

Genetic relationships among the 580 cowpea accessions were explored using two strategies. First, genetic assignment implemented in Structure version 2.3.4 (Pritchard et al., 2000) was used to identify the primary population structure in the sample. We tested *K* values from 1 to 8, with 5 replications each. The best *K* value was identified based on the Δ*K* calculation method (Evanno et al., 2005) implemented in the Structure Harvester online program (Earl and vonHoldt, 2012). We assigned genotypes to subpopulations by considering a membership coefficient (Q) value >0.75. The ancestry proportions of each sample were represented with a barplot frame in R. We estimated the spatial extent of each population cluster by projecting the genetic assignment from Structure results on a map of Africa using the tessellation approach of tess3 R package (Caye et al., 2016). Secondly, the Principal Component Analysis approach available from the R package SNPRelate (Zheng et al., 2012) was used to assess the genetic relationships between the accessions.

### *F*_ST_ Estimation

The *F*_ST_ estimator (Weir and Cockerham, 1984) implemented in the R package ‘HierFstat’ (de Meeûs and Goudet, 2007) was used to calculate *F* statistics for each SNP. We used 44,111 SNPs that passed SNP curation and 580 accessions. We considered three partitions of the dataset following the latitude. The first group included accessions with latitude > 12°N, including Egyptian and Northern Sahel accessions. The second partition was for accessions with a latitude between 0° and 12°N, including accessions from the Southern Sahara. The third group included accessions with latitude < 0° and comprises accessions from the Southern and Southeastern regions of Africa. Outliers were identified at the 99th percentile of the distribution in each comparison.

### SPA Estimation

A spatial ancestry analysis (SPA) (Yang et al., 2012) was used to identify allele frequency gradients across the sampled geographic space. The analysis used genotyping data in PLINK format and geographic coordinates (latitude and longitude) as input. No definitions of populations or sample partitions are required. We identified outliers as the top 0.1% of values.

### Environmental Genome-Wide Association Study

Two GWAS methods known for effectively managing false positives (Cebeci et al., 2023) were used to identify single-maker associations. In particular, we utilized the multi-locus GWAS models Fixed and random model circulating Probability Unification, FarmCPU (Liu et al., 2016), and the Bayesian-information Linkage Disequilibrium Iteratively Nested Keyway, BLINK (Huang et al., 2019), which are implemented in GAPIT v3 (Wang and Zhang, 2021). FarmCPU uses both the fixed effect model (FEM) and the random effect model (REM) to test markers. BLINK uses the Bayesian information content (BIC) in a FEM and linkage disequilibrium information to test markers.

We used two datasets: the first included all 580 cowpea accessions, and the second excluded 90 Southeastern accessions resulting in 490 samples. Environmental GWAS was conducted using 44,111 and 42,399 SNPs, respectively, and the 25 climatic variables described above. Quantile-quantile (Q–Q) and Manhattan plots of the P-values were used to evaluate the false positive rate. A false discovery rate (FDR) cutoff of 5% was used to identify significant associations using the R package q-value version 2.30.0 (Storey, 2022).

### Ranking of environmental variables

We used three temperature (BIO1; BIO5 and BIO8) and three precipitation-related variables (BIO12; BIO13and BIO16) to rank all georeferenced accessions relative to their ordinal value for each bioclimatic variable. We used this approach to determine putative environmental factors associated with the alternate allele’s presence in our *F_ST_* analysis. For instance, BIO 01 was reported as the average annual temperature expressed as a floating-point number at each location. This value can be sorted so that each accession takes on a discrete ordinal value regarding the rank of occurrence in our sample list. Thus, based on rank order of values per SNP, we could compare which bioclimatic variables were most associated with alternate alleles for allele frequency outliers.

### Variant annotation and Enrichment analysis of hits

Annotation for each of the iSelect variants based on the *V. uniguicalata* genome (IT97K-499-35_v1.0) (Lonardi et al., 2019) using Variant Effect Predictor (VeP) (McLaren et al., 2016) was reported by (Liang et al., 2024) and it is available at https://github.com/MorrellLAB/cowpea_annotation/tree/main/Results/IT97K-499-35_v1.0/VeP. SNPMeta (Kono et al., 2014), which performs BLAST (Camacho et al., 2009) parsing to extract annotations from the current versions of the NCBI GenBank nucleotide database (Sayers et al., 2024), was used to identify gene names and the identity of annotated copies of genes containing each of our outliers SNP in related species. Based on envGWAS, *F*_ST_ and SPA outliers, we calculated odds ratio in the functional categories (genic), coding sequence (CDS), and nonsynonymous variants. The odds ratio allows identifying if there could be more variants in than the considering regions one could expect relative to the proportions in the full genotyping data. We then used Fisher exact test to verify significance of each observe enrichment.

### Inference of the ancestral state of variants

In most cases, new adaptive functions can be attributed to derived mutations. To infer the derived versus ancestral state at each SNP, we used the maximum likelihood method (Keightley et al., 2016), updated to use multiple outgroups species and nucleotide substitution models (Keightley and Jackson, 2018) and implemented in the program est-sfs (https://sourceforge.net/projects/est-usfs/). We used genome assemblies of three *Vigna* species as outgroup sequences. These included *V. vexillata* (Zombi pea), *V. marina* (beach pea or notched cowpea), and *V. triolobata* (jungle mat bean) (Naito et al., 2022). Each of these outgroup accessions were treated as the focal species. We used phylogenetic relationships as estimated from nuclear ribosomal and chloroplast genes (Takahashi et al., 2016). We aligned the assemblies to the *V. unguicalata* genome (IT97K-499-35_v1.0) (Lonardi et al., 2019) using minimap2 v2.28 (Li, 2018) (https://github.com/MorrellLAB/cowpea_annotation). The resulting sequence alignment map (SAM) files were sorted, indexed, and converted to a binary alignment map (BAM) file using samtools v1.20 (Li et al., 2009). The alignments for each outgroup accession were converted to FASTA files using the analysis of the next generation sequencing data (ANGSD) v0.940 function ‘doFast’ (Korneliussen et al., 2014). The nucleotide state at SNP positions was extracted using the bedtools v2.31.0 function ‘getFast’ (Quinlan and Hall, 2010). Input files for ancestral state inference included a VCF of the focal species and BED files of the inferred nucleotide state. Input and output for the analysis were processed using Python scripts available at https://github.com/MorrellLAB/File_Conversions. The program est-sfs v2.04 was run using a six-rate nucleotide substitution model reported by Keightley and Jackson (2018) as R6. The program uses allele frequency information from the focal species and the nucleotide state in each outgroup species to estimate the probability that the major allele at each variant is the ancestral state. The program reports the probability that the major allele is ancestral. For probability values >50%, we reported the raw probability and the major allele. For probabilities P <50%, we reported the 1 - P and the minor allele. We reported P > 80% as the ancestral allele in an annotated VCF file. All inferred ancestral states and associated probabilities are available (along with the VCF) in a DRUM repository (https://www.dropbox.com/scl/fo/j41tc0lvkk6p5xmtrjdo8/ADn3z8diTFHVLSI98FhT42w?rlkey=jidge9a6rr32uui4o4c4vzyea&st=4a73nvmg&dl=0).

## Results

### Accession locality confirmation

The germplasm used in this study is a subset of the IITA cowpea core collection. A set of 681 accessions from Africa were initially selected, primarily categorized as landraces (632 accessions). This sample set excluded accessions where the “collection site” was a market. The accuracy of geographical information within the passport data is crucial for the success of the envGWAS approach. A detailed exploration of the passport data of the 681 accessions, including the use of the geographic sampling confirmation program ITALLIC (Onsongo et al., 2022), revealed inconsistencies in geographic origin information for 190 accessions. A detailed curation of the passport data was performed (see Methods, Fig. S2, and Table S1 for details), which resulted in a final set of 580 accessions for further analysis. The 580 accessions had been previously genotyped with 51,128 SNPs on the Illumina iSelect Cowpea Consortium Array (Muñoz-Amatriaín et al., 2017), and the genotypic data is available from Close (2023). These accessions were mapped onto the African continent with the for comparison to mean annual temperature from 1970-2000 obtained from the WorldClim2 database (Fick and Hijmans, 2017) (Fig. 1). The accessions occur across a relatively large temperature and precipitation gradient, consistent with differentiation into very distinct climatic zones (Fig. 1 & Fig. S3). Based on the current Köppen-Geiger classification (Kottek et al., 2006), most accessions in western Africa occur in the tropical savanna climatic zone (Fig. S3). Most accessions from the hottest climates in Africa occur in the tropical savanna, with a small number occurring in the hot desert climate on the southern edge of the Sahara.

**Fig. 1.**
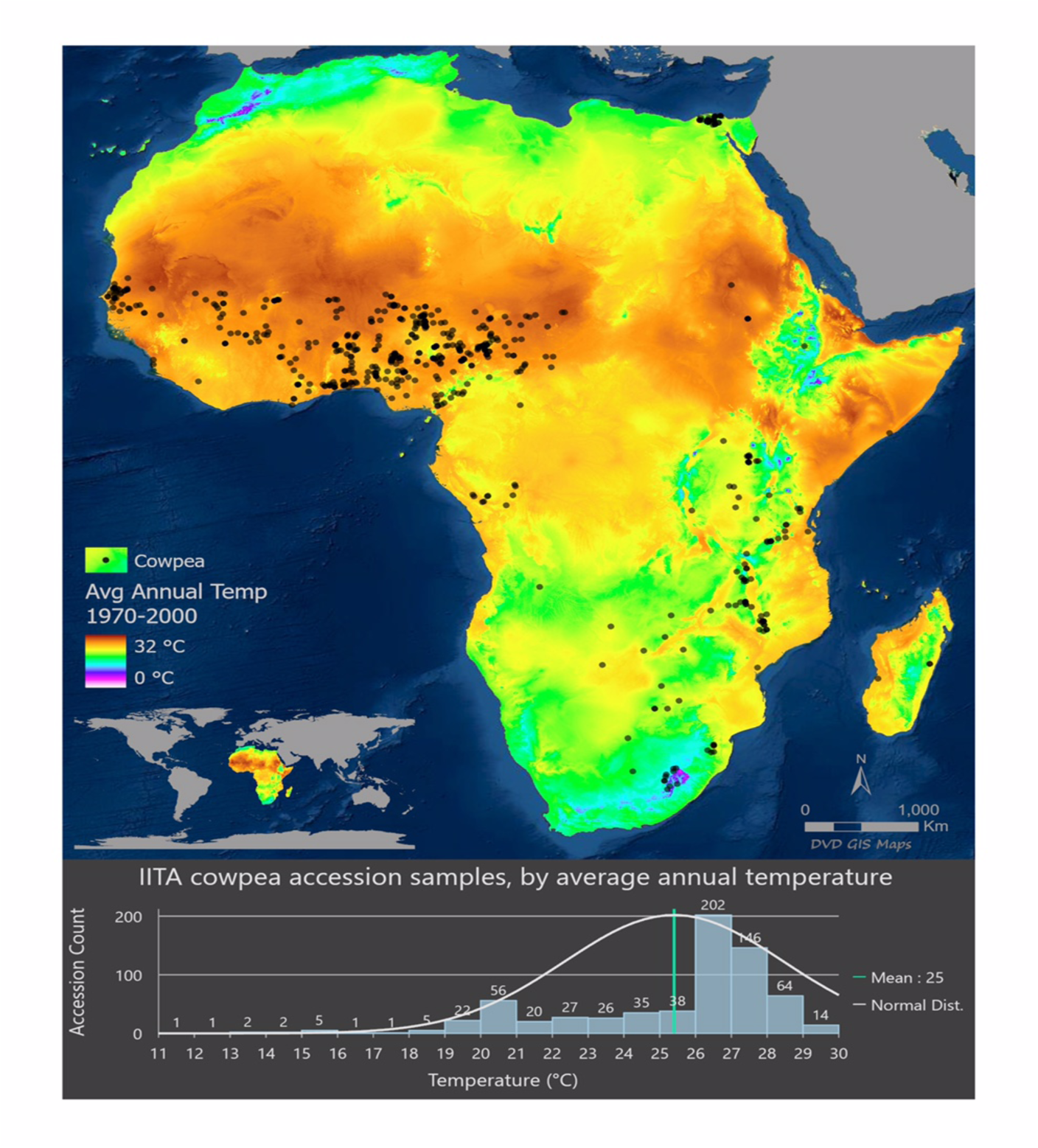
The map depicts the average mean annual temperature from 1970 to 2000. The map shows the collection site of 580 cowpea accessions as black dots. The mean annual temperature was extracted from WorldClim v2.

### Population structure

Population structure in the dataset was assessed using genetic assignment in the software Stucture (Pritchard et al., 2000) and by a principal component analysis (PCA). Using STRUCTURE outputs for a range of population numbers (*K*) between 1 and 8, Δ*K* values (Evanno et al., 2005) were calculated. Δ*K* reached a maximum at *K* = 2, which primarily distinguished Southeastern African accessions from the balance of the sample, with a secondary peak at *K* = 5 (Fig. S4). *K* = 5 was used for further analysis as it provided a more detailed description of the genetic structure in this diverse dataset (Fig. 2a). We assigned samples to populations using ancestry proportions (Q) larger than 0.75. The first genetic cluster comprised 127 accessions primarily from the African Tropical Savannah (ATS), the hottest growing region in our dataset. A second cluster included 27 accessions derived primarily from the West African Arid Steppe (WAAS), including Senegal and Gambia. A third cluster included 56 accessions from the Coastal West African Tropical area (CWAT). A fourth cluster included 105 accessions primarily from Southeastern Africa (SEA), while the fifth cluster comprised 55 accessions mostly from locations in the North African Desert (NAS), especially Egypt. The remaining 210 accessions were considered admixed (gray dots in Fig. 2b).

**Fig. 2.**
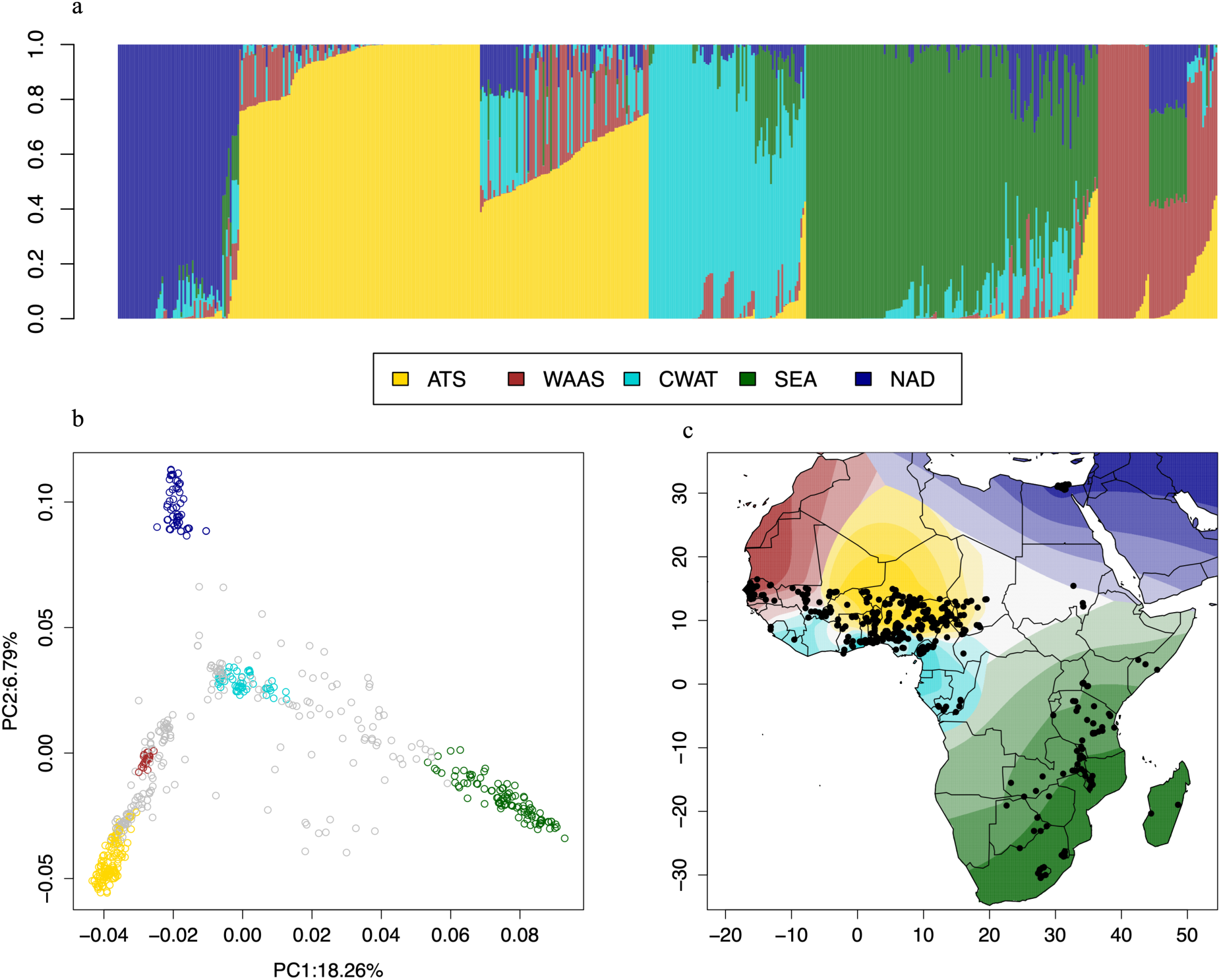
(a) Genetic assignment results for *K*=5. ATS = African Tropical Savannah (Yellow), WAAS = West African Arid Steppe (Red), CWAT = Coastal West African Tropical (Light blue), SEA = Southeastern Africa (Green), NAD = North African Desert (Dark blue). (b) Principal Component Analysis (PCA) showing genetic clustering; (c) Spatial interpolation of the geographic distribution of genetic clusters.

A principal component analysis (PCA) identified similar populations (Fig. 2b, Fig. S5). The first principal component separated samples from Egypt and Northern Africa (NAS) from samples belonging to the Southern Sahara (ATS) and Southern Africa (SEA). Accessions from western Africa (CWAT) (primarily from tropical savanna regions) and, to a lesser degree, samples from the northwestern coastal region of Africa (WAAS) are intermediate between these two extremes. The second PC separated samples from southeastern Africa (SEA) and the Southern Sahara (ATS), with other populations in an intermediate degree of differentiation. Genetic assignment based on Structure results were projected on a map of Africa using the tessellation approach implemented in Tess3 (Caye et al., 2016) (Fig. 2c).

Based on the population structure identified, with a strong genetic differentiation of the Southeastern African germplasm from the rest, which also correlates with temperature (i.e., samples from Southeastern Africa are adapted to lower growing temperatures, Fig. 1), we eliminated 90 Southeastern Africa samples for most of our further analysis and focused on the remaining 490 samples. They mainly included accessions from Western Africa, with a few samples from adjacent regions and Egypt.

### Identification of Loci Putatively Associated with Environmental Adaptation

We used measures of allele frequency-based and a mixed model association approach relative to environmental variables (envGWAS) to identify variants potentially associated with heat tolerance in cowpea. A total of 317 SNPs were identified following the workflow described in Fig. S6.

### Allele Frequency (*F*_ST_) Outliers

We examined allele frequency differentiation for three geographic partitions of cowpea accessions. We first used the sample set of 490 samples excluding Southeastern African accessions and compared samples occurring at >12°N latitude (primary in the Köppen-Geiger climate classes Bwh (hot desert) or Bsh (semi-arid climates) (Kottek et al., 2006)) that are exposed to warmer and drier growing conditions than accessions from lower latitudes in West Africa (Fig. S3). Accessions from equatorial regions are primarily from climate zone Aw (tropical wet savanna climate), with a small number of accessions from Am (tropical monsoon climate) (0°N < latitude < 12°N) (Fig. S3). This resulted in 289 of 490 accessions in this sample that occurred north of the >12°N latitude comparison. Genome-wide average *F*_ST_ = 0.039 (± 0.062), with a median of 0.011. The genetic differentiation based on *F*_ST_ was greater toward the end and lower toward the middle of each chromosome, but we did not find any genomic regions with exceptionally high *F*_ST_ values. Using an empirical cutoff of the top 0.1% of *F*_ST_ values, we identified 44 SNPs with *F*_ST_ ranging from 0.366 to 0.459 (Table 1; Table S5). The average minor allele frequency (MAF) of our *F*_ST_ outliers was 0.279 (± 0.058), consistent with the constraint on *F*_ST_ toward the potential for higher *F*_ST_ values with higher MAF.

**Table 1.** The 44 SNPs identified as outliers in *F*_ST_ analysis.

| SNP name | Chromosome | Position | SNP name | REF | ALT | MAF | $F_{ST}$ |
| --- | --- | --- | --- | --- | --- | --- | --- |
| 2_41147 | Vu01 | 35265694 | 2_41147 | T | C | 0.2691 | 0.367421642 |
| 2_23980 | Vu02 | 18984962 | 2_23980 | A | C | 0.2163 | 0.373396979 |
| 2_25308 | Vu02 | 20227677 | 2_25308 | G | T | 0.3673 | 0.371434409 |
| 2_07556 | Vu02 | 20244870 | 2_07556 | T | C | 0.2229 | 0.365564798 |
| 2_40702 | Vu02 | 2161789 | 2_40702 | T | C | 0.2327 | 0.447939419 |
| 2_25127 | Vu03 | 55523972 | 2_25127 | A | G | 0.3622 | 0.36707004 |
| 2_22916 | Vu03 | 38231022 | 2_22916 | T | C | 0.2531 | 0.368525334 |
| 2_00372 | Vu03 | 38214324 | 2_00372 | T | G | 0.3163 | 0.375015544 |
| 2_30891 | Vu04 | 14635703 | 2_30891 | A | G | 0.2163 | 0.366013666 |
| 2_40224 | Vu04 | 24273744 | 2_40224 | T | C | 0.3673 | 0.404123228 |
| 2_47878 | Vu04 | 12434298 | 2_47878 | T | C | 0.2691 | 0.383600433 |
| 2_40290 | Vu04 | 12460174 | 2_40290 | C | T | 0.2327 | 0.389740721 |
| 2_19601 | Vu05 | 3802677 | 2_19601 | G | A | 0.2531 | 0.407273132 |
| 2_18508 | Vu05 | 3708611 | 2_18508 | C | T | 0.2229 | 0.368754761 |
| 2_53903 | Vu05 | 3739351 | 2_53903 | A | C | 0.3163 | 0.404886057 |
| 2_25041 | Vu05 | 3835894 | 2_25041 | G | T | 0.3622 | 0.430975934 |
| 2_27980 | Vu05 | 5936454 | 2_27980 | G | T | 0.2691 | 0.365169439 |
| 2_08871 | Vu05 | 5942506 | 2_08871 | T | C | 0.2327 | 0.392285478 |
| 2_28186 | Vu07 | 26109238 | 2_28186 | C | T | 0.2163 | 0.388022533 |
| 2_21861 | Vu07 | 26137988 | 2_21861 | A | G | 0.3673 | 0.388022533 |
| 2_48714 | Vu07 | 26149694 | 2_48714 | C | T | 0.2229 | 0.38890675 |
| 2_49476 | Vu07 | 26156897 | 2_49476 | T | C | 0.3163 | 0.388022533 |
| 2_25770 | Vu07 | 26168611 | 2_25770 | C | T | 0.2531 | 0.440645803 |
| 2_03060 | Vu07 | 26173924 | 2_03060 | A | G | 0.3622 | 0.442324938 |
| 2_00885 | Vu08 | 34707621 | 2_00885 | A | C | 0.2327 | 0.404014544 |
| 2_07072 | Vu08 | 30038329 | 2_07072 | C | T | 0.3622 | 0.369921954 |
| 2_00857 | Vu08 | 7392202 | 2_00857 | T | G | 0.2691 | 0.366142054 |
| 2_09973 | Vu08 | 7405223 | 2_09973 | C | A | 0.2327 | 0.366142054 |
| 2_29223 | Vu08 | 7406627 | 2_29223 | C | T | 0.2163 | 0.372196772 |
| 2_51987 | Vu08 | 7408089 | 2_51987 | C | T | 0.3673 | 0.372196772 |
| 2_53354 | Vu08 | 7412329 | 2_53354 | T | C | 0.2229 | 0.372872684 |
| 2_51407 | Vu08 | 7419005 | 2_51407 | C | T | 0.2531 | 0.366446259 |
| 2_52823 | Vu08 | 7417594 | 2_52823 | A | G | 0.3163 | 0.378947462 |
| 2_06639 | Vu08 | 30099911 | 2_06639 | A | G | 0.2691 | 0.365935158 |
| 2_51380 | Vu09 | 38945148 | 2_51380 | C | T | 0.3673 | 0.451429091 |
| 2_05643 | Vu09 | 41247766 | 2_05643 | A | G | 0.3163 | 0.384307282 |
| 2_25481 | Vu09 | 41181696 | 2_25481 | C | T | 0.2229 | 0.384554079 |
| 2_14464 | Vu09 | 41270428 | 2_14464 | C | A | 0.2531 | 0.386258619 |
| 2_14668 | Vu09 | 31846740 | 2_14668 | T | C | 0.2163 | 0.37395871 |
| 2_19641 | Vu11 | 39324907 | 2_19641 | C | A | 0.3673 | 0.428637588 |
| 2_25564 | Vu11 | 35441960 | 2_25564 | G | A | 0.3622 | 0.458509445 |
| 2_17698 | Vu11 | 39316305 | 2_17698 | T | C | 0.2163 | 0.398862745 |
| 2_02052 | Vu11 | 35816995 | 2_02052 | G | T | 0.2691 | 0.39222921 |
| 2_02051 | Vu11 | 35817750 | 2_02051 | C | A | 0.2327 | 0.39222921 |

The highest *F*_ST_ values included 10 SNPs on chromosome Vu08; seven occurred within a 27-kb genomic region that includes two genes (*Vigun08g058300* and *Vigun08g058400).* The second largest block of high *F*_ST_ variants included six SNPs on a 65 kb region of chromosome Vu07. The first of these six SNPs, from the 5’ end of the region was SNP 2_28186, where the alternate and derived allele was found most abundant in samples from the western Sahel region (Fig. 3c). Near the beginning of the region (∼26,113,000 bp), two gene models encoding DNAJ heat shock proteins, *Vigun07g150800* and *Vigun07g150900,* are located (Fig. 3a). DNAJ has been described as a heat shock protein playing a key role in the growth and the response of capsicum (*Capsicum annuum* L.) to heat stress (Fan et al., 2017).

**Fig. 3.**
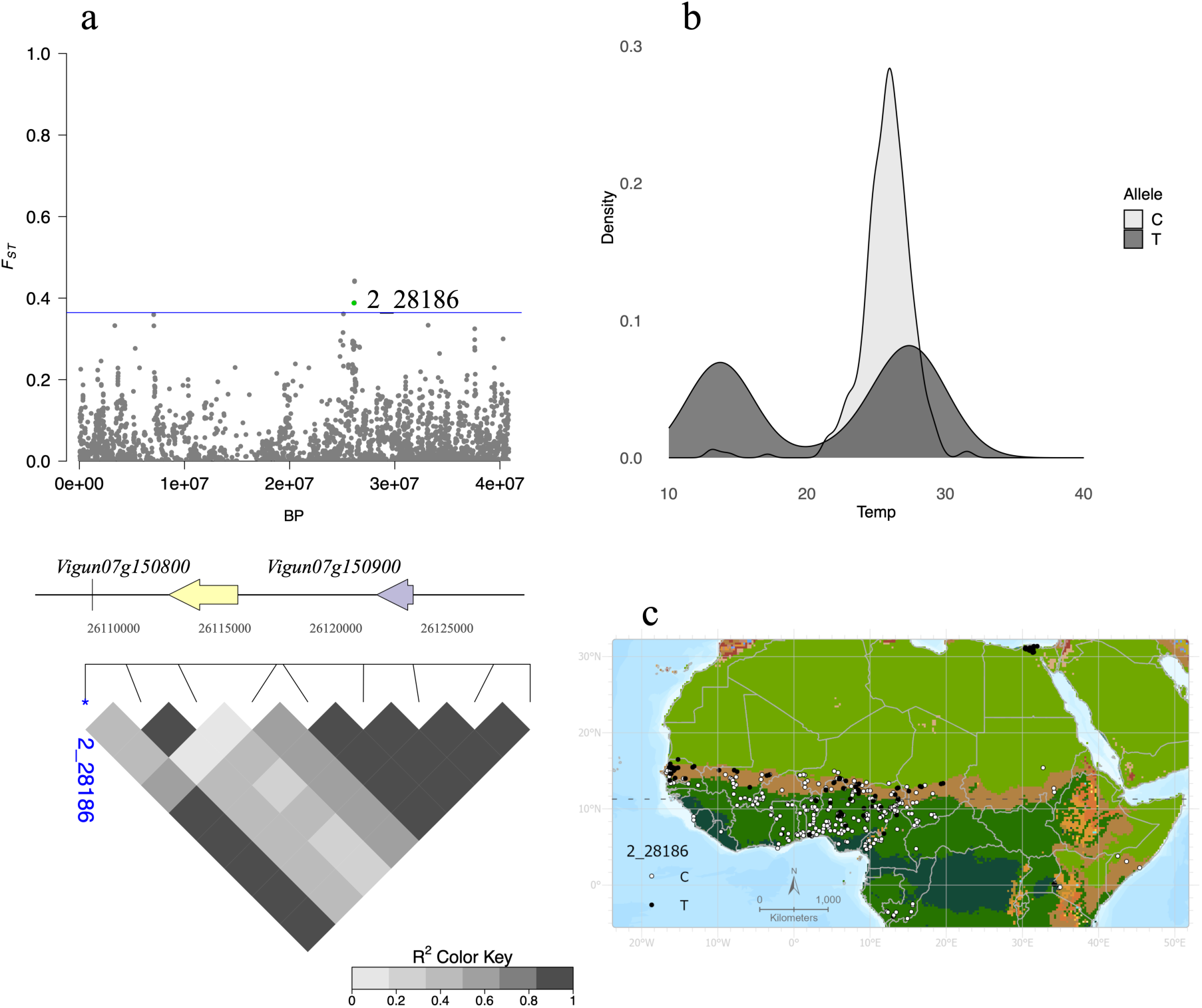
(a) *F*_ST_ values for Vu07 are depicted in a Manhattan plot. The two genes in closest proximity to SNP 2_28186 are depicted, along with pairwise LD for nine SNPs in their proximity. (b) The distribution of 2_28186 alleles across a temperature range. (c) The geographic distribution of alleles at 2_28186 in West Africa.

With an average MAF of 0.29, the alternate alleles for these six variants (2_28186, 2_21861, 2_48714, 2_49476, 2_25770, 2_03060) occur in accessions growing across a broad geographic range that also includes the hottest localities. Using a series of nine *F*_ST_ outliers, we identified relatively high levels of linkage disequilibrium in that ∼65 kb region, with an average *r*^2^ of 0.62 (Fig.3.a).

*F*_ST_ comparisons between samples from the African equatorial region (0°N < latitude < 12°N) and those from Southern Africa (latitude < 0°N) resulted in much higher values (mean *F*_ST_ = 0.231 ± 0.180, median = 0.201). A comparison of populations from higher latitudes (>12°N) to Southern African accessions resulted in *F*_ST_ = 0.197 (±0.149) with a median of 0.181 a (Fig. 3a & b). The demographic history of dissemination of cultivated cowpea has not been extensively explored. Still, elevated *F*_ST_ values for populations from Southern Africa, could suggest very different demographic histories for cultigens from various portions of the species’ range in Africa.

### Allele Frequency Gradients (SPA)

Using the same 0.01% empirical cutoff for 42,339 SNPs, the SPA analysis yielded 42 outliers with SPA values > 6.227 (Table 2). SPA values ranged from 0.002 - 6.725 with a median of 1.814 and a mean of 2.123 (± 1.459). The largest SPA values were observed for two SNPs (2_02051 and 2_02052) on Vu11, which were also identified as outliers in the *F*_ST_ analysis (Fig. 4a). The two SNPs followed a nearly identical geographic pattern, with the alternate (and ancestral) alleles most abundant in West Africa and becoming less common in cooler, wetter climates to the southeast (Fig. S6). Both SNPs occurred in introns of the gene model *Vigun11g148700* (Fig. 4a), annotated as an ethylene-responsive transcription factor, which belongs to a gene family previously associated with heat stress response in *Arabidopsis thaliana* (Huang et al., 2021). High LD values were also observed in that same region on Vu11. A series of 14 SPA outliers on chromosome Vu11 spanned a region of ∼106 kb with a mean *r*^2^ value of 0.44 (Fig. 4a). But the largest group of SPA outliers occurred in 11 of the 12 SNPs, on a Vu08 region spanning 352 kb. This region includes 50 genes and does not overlap with variants found by envGWAS or *F*_ST_ outlier analyses.

**Fig. 4.**
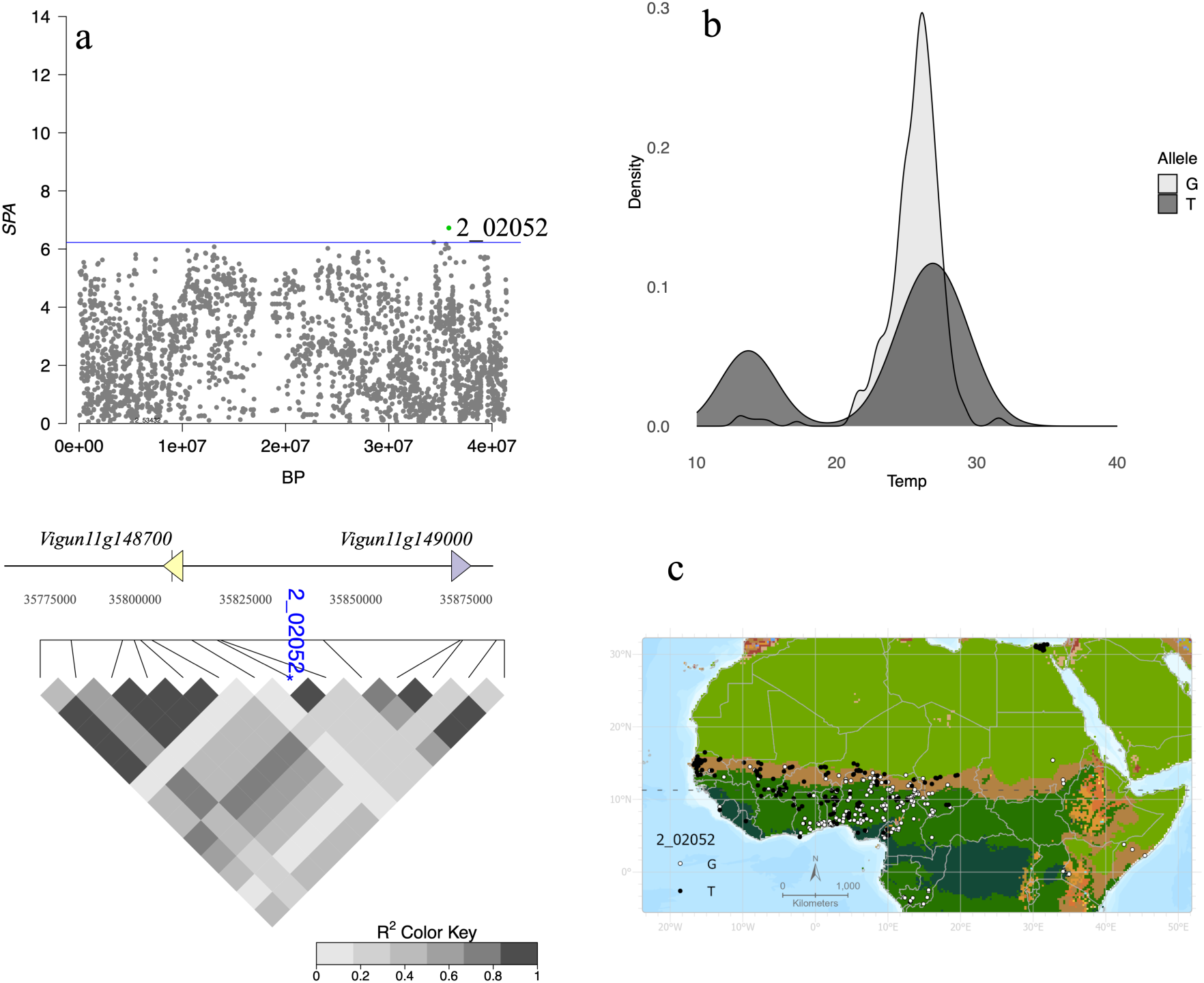
(a) SPA values for Vu11 are depicted in a Manhattan plot. The 2 genes in closest proximity to the SNP 2_02052 are depicted along with pairwise LD for 14 SNPs in the proximity. (b) The distribution of alleles at 2_02052 is the highest value of SPA outlier. (c) Geographic distribution of alleles at 2_02052 in Western African cowpea accessions.

**Table 2.** The 42 SNPs identified as outliers in SPA analysis.

| Chromosome | Position | SNP | REF | ALT | MAF | SPA |
| --- | --- | --- | --- | --- | --- | --- |
| Vu02 | 9772009 | 2_40911 | C | T | 0.119388 | 6.33218595 |
| Vu02 | 13201161 | 2_41585 | C | T | 0.131633 | 6.34684379 |
| Vu02 | 32217978 | 2_15286 | C | T | 0.130612 | 6.38583729 |
| Vu02 | 33609908 | 2_31898 | G | T | 0.119388 | 6.26171032 |
| Vu02 | 33732878 | 2_22825 | T | G | 0.138776 | 6.26811305 |
| Vu04 | 5583939 | 2_42020 | C | T | 0.118896 | 6.53341899 |
| Vu04 | 12460174 | 2_40290 | C | T | 0.718367 | 6.30143805 |
| Vu04 | 24273744 | 2_40224 | T | C | 0.158436 | 6.30983993 |
| Vu05 | 366583 | 2_11664 | T | C | 0.152041 | 6.48889183 |
| Vu05 | 371882 | 2_30578 | G | A | 0.156122 | 6.4797115 |
| Vu05 | 618953 | 2_06865 | C | T | 0.126531 | 6.46012734 |
| Vu05 | 761491 | 2_49015 | G | A | 0.127551 | 6.44281203 |
| Vu05 | 1373737 | 2_54794 | C | T | 0.189331 | 6.28440948 |
| Vu05 | 3728035 | 2_16071 | T | C | 0.130612 | 6.30292467 |
| Vu05 | 3738124 | 2_05074 | A | G | 0.121429 | 6.361751 |
| Vu05 | 12163327 | 2_07002 | G | T | 0.403974 | 6.31524117 |
| Vu05 | 46120114 | 2_25170 | G | A | 0.115942 | 6.43666647 |
| Vu05 | 47493777 | 2_18368 | A | G | 0.134969 | 6.43587753 |
| Vu07 | 3369363 | 2_30853 | G | T | 0.12449 | 6.62377586 |
| Vu07 | 5310677 | 2_02754 | A | G | 0.126294 | 6.25800453 |
| Vu07 | 5721012 | 2_46140 | T | C | 0.137149 | 6.42867755 |
| Vu07 | 26168611 | 2_25770 | C | T | 0.222904 | 6.3098184 |
| Vu07 | 26173924 | 2_03060 | A | G | 0.216327 | 6.35519846 |
| Vu08 | 255408 | 2_16556 | A | G | 0.129592 | 6.34855549 |
| Vu08 | 263699 | 1_0684 | C | A | 0.126531 | 6.49674083 |
| Vu08 | 273444 | 2_29455 | A | G | 0.152083 | 6.2905076 |
| Vu08 | 292459 | 2_23087 | T | G | 0.129592 | 6.25917809 |
| Vu08 | 394592 | 2_01893 | G | A | 0.135714 | 6.36260038 |
| Vu08 | 410767 | 2_15263 | C | A | 0.139706 | 6.33953461 |
| Vu08 | 411790 | 2_15262 | C | T | 0.135714 | 6.34272106 |
| Vu08 | 416063 | 2_33384 | A | T | 0.156122 | 6.23319836 |
| Vu08 | 416227 | 2_31016 | A | C | 0.156122 | 6.23319836 |
| Vu08 | 420378 | 2_04503 | G | T | 0.161224 | 6.22784024 |
| Vu08 | 607696 | 2_09210 | T | C | 0.127551 | 6.27960781 |
| Vu08 | 27486972 | 2_32541 | C | G | 0.118367 | 6.53221898 |
| Vu09 | 29038851 | 2_14338 | C | T | 0.130612 | 6.33701787 |
| Vu09 | 32114933 | 2_15819 | T | G | 0.877551 | 6.37556006 |
| Vu09 | 34064769 | 2_26718 | A | G | 0.881633 | 6.33974334 |
| Vu10 | 37534007 | 2_15395 | C | T | 0.12449 | 6.36096369 |
| Vu11 | 34351838 | 2_50899 | C | T | 0.169388 | 6.23215601 |
| Vu11 | 35816995 | 2_02052 | G | T | 0.383673 | 6.72527267 |
| Vu11 | 35817750 | 2_02051 | C | A | 0.383673 | 6.72527267 |

For SPA outliers, the potential nature of selective factors that could contribute to steep allele frequency gradients is less evident because, unlike *F*_ST_ comparisons, the observed allele frequency differentiation occurred in many different geographic partitions of the sample. Comparing SPA outliers to temperature and precipitation environmental variables at the localities that carry the minor allele identified, for most of the SPA outliers, annual precipitation (BIO12) had the lowest rank values, suggesting that precipitation drives the allele frequency gradients (Fig. S7 and S8). The only exceptions were three SNPs (2_02051, 2_02052, and 2_07002) where the maximum temperature of the warmest month (BIO5) showed the lowest rank (Table S4).

### Environmental Genome-Wide Association Study

We used the multi-locus mixed models FarmCPU and BLINK implemented in the R package GAPIT3 to investigate 25 climatic variables for the environmental association. We used an FDR threshold of 5% to detect significant associations. As bioclimatic variables are highly correlated (Fig. S1), we focused only on significant SNPs associated with at least two variables of temperature or precipitation. Using 490 samples excluding those from southeastern Africa, we identified 89 SNPs associated with temperature and precipitation variables. When using all 580 accessions in the dataset, we detected 149 SNPs associated either with temperature (TEMP) variables (BIO1 to BIO11 + IC3, mean MAF = 0.131 ±0.115) or precipitation (PREC) variables (BIO12 to BIO19 + IC1 + IC2, mean MAF = 0.134 ±0.132) (Table S7). We summarized the variables identified into five groups. The TEMP group includes 87 SNPs associated with BIO1 - BIO11 or IC3. The PREC group includes149 SNPs associated with BIO12 - BIO19 or IC1 and IC2. The HIGH group has 11 SNPs associated with BIO5, BIO9, or BIO10. The DRY group contains 24 SNPs associated with eitherBIO14, BIO17, or BIO18. Finally, the HEAT group includes 11 SNPs associated with TMIN or TMAX. Altogether, we identified 219 unique SNPs (i.e., 19 SNPs were detected in both datasets, and some of them were associated with more than one variable) (Table S5).

For TEMP hits, some genomic regions stood out, with individual SNPs distributed on chromosomes Vu04, Vu05, Vu07, Vu10, and Vu11 (MAF<5%). These hits were also associated with the PREC variables and can be useful in cowpea breeding for tolerance to high temperatures.

### Environmental variables ranking

Allele frequency differentiation identified loci that differed in variant frequency between regions but provided limited information about the potential nature of differences between localities that could induce selective changes in allele frequency. We used three temperature and three precipitation-related bioclimatic factors that can be ranked from most to least extreme value to determine if these factors could explain allele frequency differences at outlier loci. Temperature values for accessions were sorted such that the highest temperatures or lowest precipitations have the smallest values (highest ranks). For samples with elevated *F*_ST_ values, the maximum temperature of the warmest month (BIO5) showed the lowest rank for all outliers. The second lowest-ranked variable was the mean temperature of the wettest quarter (BIO8).

### Overlap of variants detected

We found extensive overlap for variants identified in envGWAS associated with TEMP and PREC, as depicted in an upset plot (Fig. 5). We found no overlap between variants identified in the envGWAS and *F*_ST_ analysis or SPA results described above. This may result partly from very different MAF for *F*_ST_ and SPA outliers and associations found with envGWAS.

**Fig. 5.**
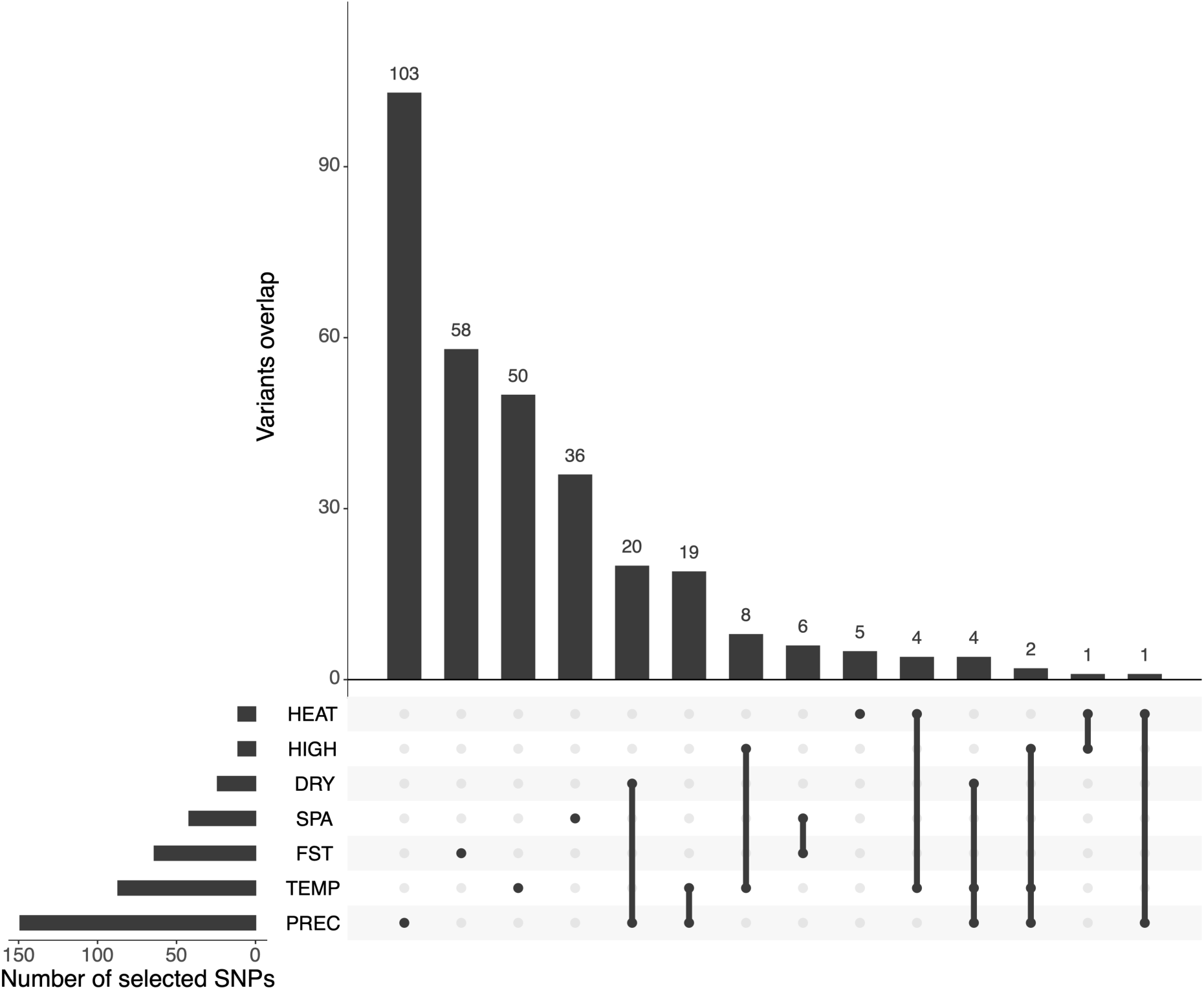
The overlap of variants between different types of climatic variables and allele frequency approaches. There is considerable overlap of SNP markers, occurring at both high and low frequencies in the population. Horizontal bars show the number of selected SNP per type of climatic variables; vertical bars present the size of the overlap of variants. Black dot indicates the types of climatic variables which overlapping size is represented in the vertical bar.

The *F*_ST_ and SPA results produced some overlapping variants. The highest absolute values for SPA involved SNPs 2_02051 and 2_02052 (Table 2), located within the same single gene (*Vigun11g148700)* on Vu11with these same SNPs also found as *F*_ST_ outliers. Additional SNPs overlapping between *F*_ST_ and SPA were found on Vu04 (2_40290 and 2_40224) and Vu07 (2_03060 and 2_25770). Two SPA outliers on Vu05 (2_05074 and 2_16071) occurred in a region with elevated *F*_ST_ values, but the same variants were not involved.

### Variant annotation and Enrichment analysis of hits in three functional categories

BLAST searches of Genbank database, based on the contextual sequence around each outlier SNP, identified matches in the National Center for Biotechnology Information (NCBI) GenBank nucleotide database (Sayers et al., 2024) for 150 of 317 variants identified in our various comparative analyses. This analysis was implemented in SNPMeta (Kono et al., 2014). A table of SNPMeta output for each of the SNPs detected as outliers in our analyses is reported in Table S7. Among the variants producing BLAST results were 2 SNPs found within heat shock proteins (1_0346 and 2_05343) and 3 SNPs found within various cytochrome P450 gene family members (2_21837, 2_33252, 2_246982).

We performed an enrichment analysis calculating an odds ratio to determine if there were proportionally more variants in functional categories (genic), coding sequence (CDS), and nonsynonymous than we would expect relative to the proportions in the full genotyping data. Our 317 candidate outliers from envGWAS, *F*_ST_ and SPA were enriched for SNPs in genic regions (1.11), for CDS (1.21) and for nonsynonymous SNPs (1.31). However, the observed trend was non-significant in a Fisher exact test, with the greatest enrichment for CDS resulting in p = 0.08 (Table 3).

**Table 3.** Enrichment of candidate SNPs within functional categories.

| Category | SNPs in candidate | SNPs out candidate | SNPs in full data | SNPs out full data | Odds Ratio | P-value |
| --- | --- | --- | --- | --- | --- | --- |
| Genic | 207 | 110 | 27710 | 16396 | 1.11 | 0.20 |
| CDS | 83 | 234 | 9977 | 34129 | 1.21 | 0.08 |
| Nonsynonymous | 39 | 278 | 4272 | 39834 | 1.31 | 0.07 |

SNPs in candidate = Number of SNPs in the category in the candidate list
SNPs out candidate = Number of SNPs not in the category in the candidate list
SNPs in full data = Number of SNPs in the category in the full data SNPs out full data = Number of SNPs not in the category in the whole dataset Odds Ratio = (SNPs in candidate x SNPs out full data) / (SNPs out candidate x SNPs in full data)

### Ancestral state inference

Alignments of outgroup samples to the cowpea reference genome produced the most complete alignments based on the minimap2 setting “ASM20”, or divergence of not more than 10%. The nucleotide states for *V. unguiculata* iSelect SNPs were identified by querying the assemblies of three cowpea relatives aligned to the cowpea reference genome. This resulted in the identification of nucleotide state for 12,869 SNPs for *V. vexillata*; 10,955 for *V. trilobata* and 10,281 for *V. marina*. Using estSFS (Keightley and Jackson, 2018) for maximum-likelihood inference of ancestral state, 29,218 SNPs were identified with an inferred probability of correct inference of ancestral states of >80%. The alternate allele was determined to be ancestral for 15,322 SNPs, with the reference allele ancestral for 13,896 SNPs. The ancestral state of each selected SNP is indicated in Table 2 (*F_ST_* outliers), Table 3 (SPA outliers) and Table S5 (envGWAS outliers).

## Discussion

Based on envGWAS, we identified 98 variants that survived FDR correction for association with heat-related variables. There were significant associations on all chromosomes, mostly detected on chromosomes Vu03 (13 SNP) and Vu04 (12 SNP). Both *F*_ST_ and SPA analysis identified variants that are also putatively associated with adaptive differences across the species range. The comparison of alternate allelic states to environmental variables shows that all the *F*_ST_ were most strongly associated with temperature differentiation, while SPA outliers were associated with precipitation differences. This difference reflects the way outliers are defined in these two approaches. By delineating samples on either side of 12°N latitude, we divided cowpea lines from hot semi-arid climates and hot desert regions in the southern Sahara from those from the tropical savannah, essentially forcing outlier allele frequency to occur as differences in temperature. This has the desirable effect of identifying variants that occur more frequently in lines from hotter environments. However, because SPA identifies gradients in allele frequency without arbitrary partitions of the sample, it could be argued that differentiation in allele frequency identified may be more closely associated with selection imposed by the environment in shaping allele frequencies. That is, selection for differences in precipitation may have been the more important adaptive force in cowpea. This observation appears consistent with a comprehensive environmental association study in 25 plant species that concludes that precipitation-related variation is more frequently encountered than variants associated with temperature difference (Whiting et al., 2024). This may occur in part because changes in temperature have occurred more recently, resulting in fewer generations of selective pressure on adaptation to increasing temperature within environments (Whiting et al., 2024).

We identified a much larger number of variants significantly associated with precipitation-related environmental variables than with heat-related variables. This resulted in 72 of the 149 precipitation-related variants also detected as outliers for heat-related variables (Fig. 6). Prima facie, this suggested that precipitation differences across the geographic range of our sample were important determinants of allelic differentiation among samples, and perhaps also that these traits were inextricably linked. While it is difficult to make inferences about the physiological basis of heat stress based on these results, many variants with the strongest association with heat stress are also associated with water availability (Fig. 6).

As noted above, envGWAS and the allele frequency outlier approaches identified largely independent sets of variants. However, *F*_ST_ and SPA results overlapped, particularly for some extreme allele frequency outliers. At times, a lack of overlap among the variants detected in these various approaches has led to speculation regarding the veracity of the results. For example, Anderson et al. (2016) found little overlap between these same comparisons, i.e., envGWAS, *F*_ST_, and SPA, in a study in *Glycine soja*, the closest wild relative of cultivated soybean (*Glycine max*). However, Lei et al. (2019) in seeking to determine if previously cloned genes involved in environmental differentiation in barley landrace could be detected in envGWAS and allele frequency comparisons, found many well-characterized genes identified as significant based on envGWAS, one of several *F*_ST_ comparisons or both. There are several well-characterized genes with known effects in barley that were found in multiple comparisons. For example, Phytochrome C, a flowering time-related locus, was identified using four bioclimatic variables and an elevation *F*_ST_ comparison (Lei et al., 2019). Other well-characterized genes were identified using only one of the approaches, suggesting that the approaches are at times complementary (Campbell et al., 2025). These results may also reflect some of the challenges of detection of variants based on factors such as SNP density (Lei et al., 2019) and the complexity of environmental factors that can drive adaptation (Lotterhos, 2023).

Approaches for improving the detection of variants associated with adaptation to the environment are being actively explored (Campbell et al., 2025). One approach exploits linkage disequilibrium among linked variants (Booker et al., 2024) to increase the detection of adaptive variants. The conditions in which this approach improves detection partially dependent on levels of variation, recombination rates, and marker density. Another recently proposed approach uses inference of geographic provenance based on genotype composition (Battey et al., 2020) [Battey et al. 2020] to create a much denser grid of sampled individuals, including samples that did not originally include detailed location of origin information (Chang and Schmid, 2025).

There is considerable potential to use our results to identify cowpea genetic resources likely to carry variants relevant to increased heat tolerance in cowpea lines grown as a critical food source in regions where heat stress is likely to become increasingly important. This process could begin with a focus on variants detected here that occurred at a relatively low frequency among current breeding materials from programs in West Africa (Muñoz-Amatriaín et al., 2017) and extend to machine-learning based methods that enable the annotation of variants beyond coding regions (Benegas et al., 2023; Zhai et al., 2024).

## Acknowledgments

This study was funded by the Foundation for Food & Agriculture Research (Award# ICRC20-0000000032) to EFR, KJB, MA-M, OB, and PLM. The authors thank Tchamba Marimagne (IITA Genebank, Nigeria) for his valuable input during passport data curation, and Fiona Todd (University of Minnesota, USA) for the curation of materials associated with the manuscript. This research was carried out with software and hardware support provided by the Minnesota Supercomputing Institute (MSI) at the University of Minnesota. This research benefited from the advice and guidance of Timothy J. Close (U. of California Riverside, USA).

## Author Contributions

RA, EJL, JBP, KMV, KMV, HA, MA-M and PLM performed data analysis. RA, JBP, KMV and PLM wrote the manuscript. RA, MA-M, EJL, MKB, LG, EFR, OB, KJB, and PLM designed the research. EFR, KJB, MA-M, OB, and PLM acquired funding. All authors reviewed and edited the manuscript.

## Data Availability

Genotyping data was derived from the cowpea Illumina iSelect genotype platform and was previously reported (Fiscus et al., 2024) and can be downloaded from (https://doi.org/10.6086/D19Q37). Genotyping data used in this study is available in PLINK and VCF format, using updated physical positions of variants as reported by Liang et al. (2024). Files are available from a Data Repository for University of Minnesota (DRUM) repository (currently at https://www.dropbox.com/scl/fo/j41tc0lvkk6p5xmtrjdo8/ADn3z8diTFHVLSI98FhT42w?rlkey=jidge9a6rr32uui4o4c4vzyea&st=4a73nvmg&dl=0; DOI to be added after review). A Keyhole Markup Language (KML) file is provided for sample localities and can be used to display localities using Google Earth. All code used for data analysis and figure generation is available from https://github.com/MorrellLAB/cowpea_environmental

## Supporting information

https://www.dropbox.com/scl/fo/u6645n1q3p8lxgrebujp6/AJ1lv8u6jTOKNW2yQloQMOY?e=2&preview=Table1.csv&rlkey=s71wd27c00mu6jt8wuf2isofl&dl=0

## Supporting Information

Table S1. A list of no curated, genotyped accessions

Supplemental file

Table S2. The list of 580 accessions used in this study

Supplemental file

**Table S3.** The 22 environmental variables used in this study.

| Variable | Meaning |
| --- | --- |
| BIO1 | Annual Mean Temperature |
| BIO2 | Mean Diurnal Range (Mean of monthly (max temp - min temp)) |
| BIO3 | Isothermality (BIO2/BIO7) ($\times 100$) |
| BIO4 | Temperature Seasonality (standard deviation $\times 100$) |
| BIO5 | Max Temperature of Warmest Month |
| BIO6 | Min Temperature of Coldest Month |
| BIO7 | Temperature Annual Range (BIO5-BIO6) |
| BIO8 | Mean Temperature of Wettest Quarter |
| BIO9 | Mean Temperature of Driest Quarter |
| BIO10 | Mean Temperature of Warmest Quarter |
| BIO11 | Mean Temperature of Coldest Quarter |
| BIO12 | Annual Precipitation |
| BIO13 | Precipitation of Wettest Month |
| BIO14 | Precipitation of Driest Month |
| BIO15 | Precipitation Seasonality (Coefficient of Variation) |
| BIO16 | Precipitation of Wettest Quarter |
| BIO17 | Precipitation of Driest Quarter |
| BIO18 | Precipitation of Warmest Quarter |
| BIO19 | Precipitation of Coldest Quarter |
| TMIN | Minimum Temperature of the approximate flowering Period |
| TMAX | Maximum Temperature of the approximate flowering Period |
| PREC | Precipitation of the approximate flowering Period |

**Table S4.** Bioclim coordinate on three first Independent Components. We used the first three ICs that explained 81.18% of the bioclimatic data information. The IC1 summarize the precipitation of the wettest month, IC2 explained the precipitation of the driest period, and IC3 represented the global temperature data.

| Bioclim | IC1 | IC2 | IC3 |
| --- | --- | --- | --- |
| BIO1 | -0.391 | 0.407 | -0.809 |
| BIO2 | 0.228 | 0.853 | 0.194 |
| BIO3 | -0.718 | -0.371 | -0.176 |
| BIO4 | 0.837 | 0.217 | 0.302 |
| BIO5 | -0.036 | 0.73 | -0.629 |
| BIO6 | -0.607 | -0.176 | -0.755 |
| BIO7 | 0.557 | 0.733 | 0.241 |
| BIO8 | -0.665 | 0.46 | -0.406 |
| BIO9 | 0.064 | -0.088 | -0.934 |
| BIO10 | -0.048 | 0.563 | -0.771 |
| BIO11 | -0.606 | 0.239 | -0.742 |
| BIO12 | -0.846 | -0.411 | -0.034 |
| BIO13 | -0.888 | 0.013 | -0.076 |
| BIO14 | -0.426 | -0.566 | 0.229 |
| BIO15 | 0.062 | 0.902 | -0.11 |
| BIO16 | -0.883 | -0.052 | -0.038 |
| BIO17 | -0.445 | -0.616 | 0.157 |
| BIO18 | -0.662 | -0.391 | 0.39 |
| BIO19 | -0.433 | -0.452 | -0.317 |

Table S5. List of envGWAS outliers

Supplemental file

Table S6. Extreme allele in *F*ST

Supplemental file

Table S7. SNP Meta output

Supplemental file

**Fig. S1.**
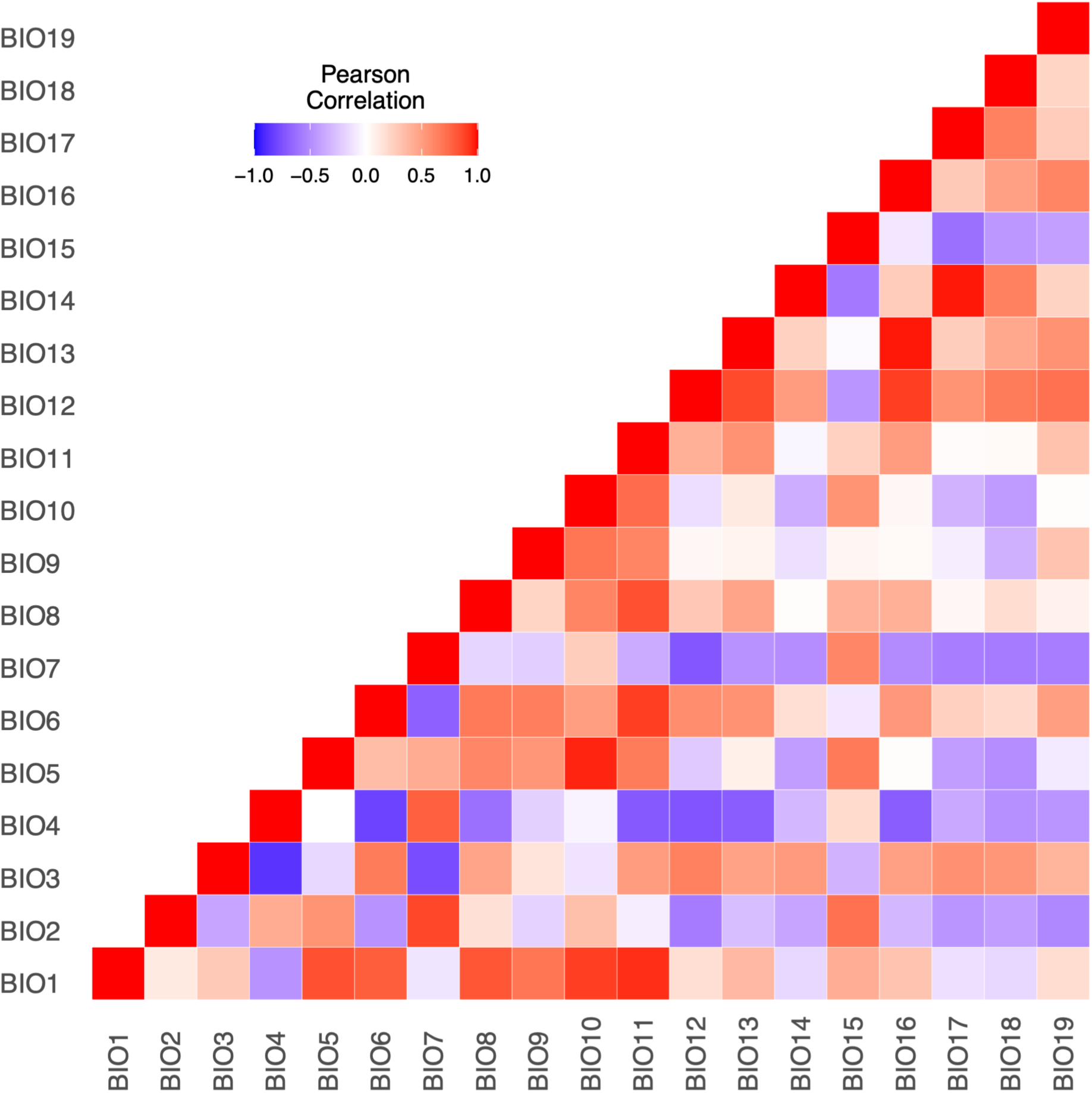
Pearson pairwise correlation matrix of Bioclim variables. The heatmap represents correlation coefficients between the 19 Bioclim variables. The red indicating strong positive correlations and blue indicating strong negative correlations. The color gradient ranges from - 1.0 (strong negative correlation) to 1.0 (strong positive correlation.

**Fig. S2.**
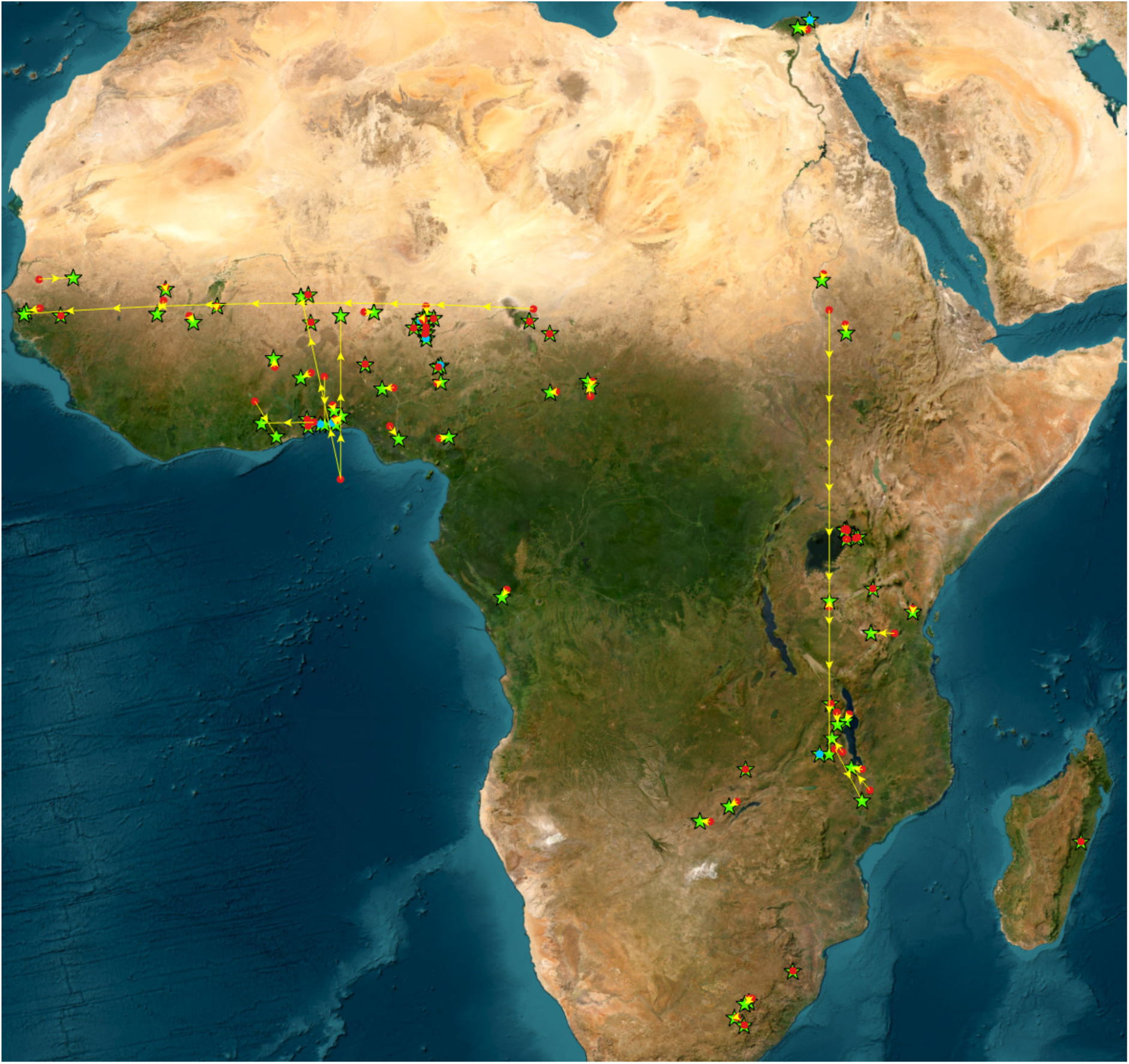
Location verification results for cowpea accessions. Red points indicate geographic coordinates from the original passport file, while the green points show the verified or updated geolocation for the accession. Yellow lines show paths between original and updated locations.

**Fig. S3.**
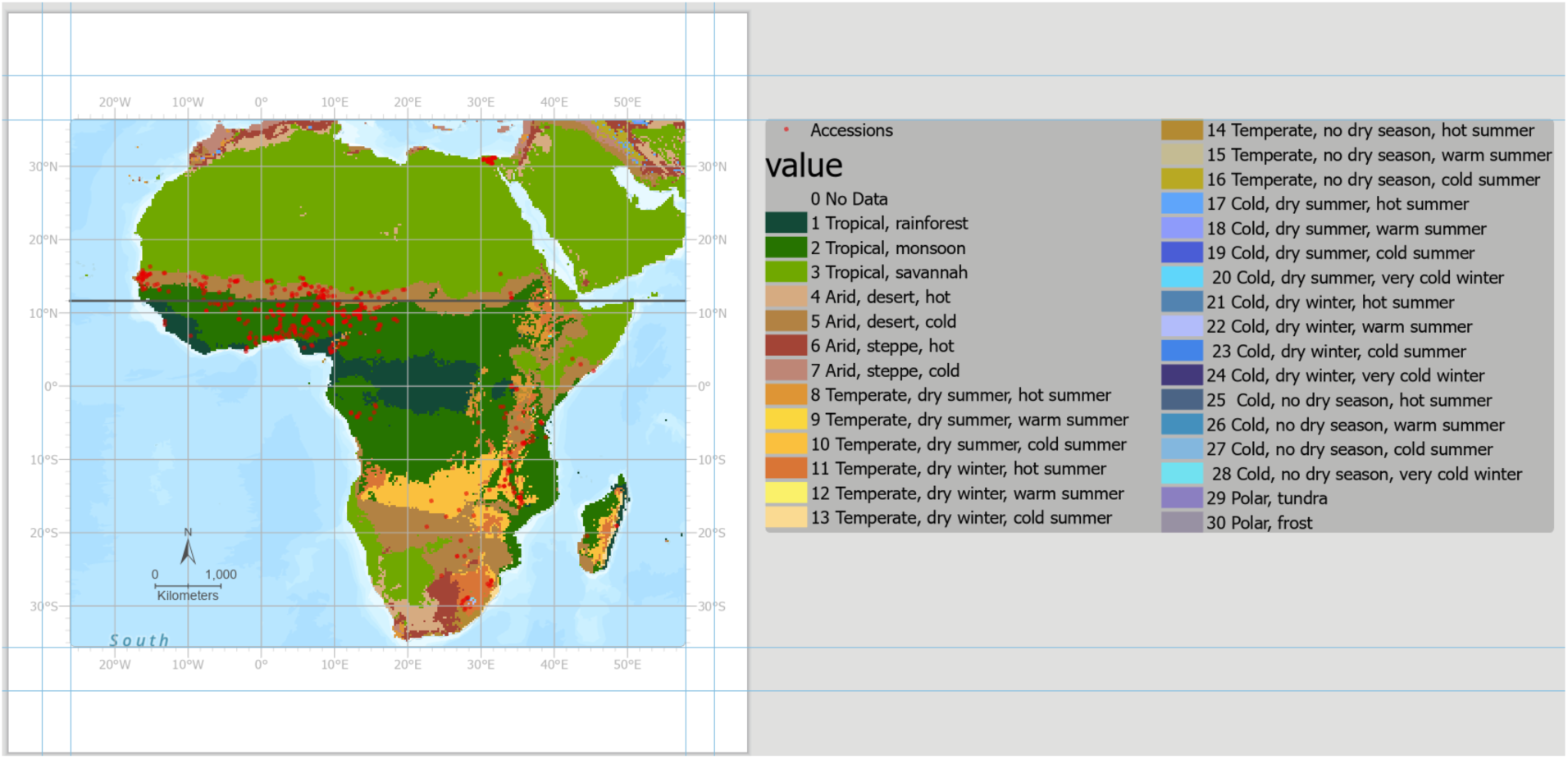
A depiction of the Köppen-Geiger climatic zone across the African continent. The collection locality of cowpea accessions is shown. The line at 12°N latitude indicates the partition used for allele frequency comparisons.

**Fig. S4.**
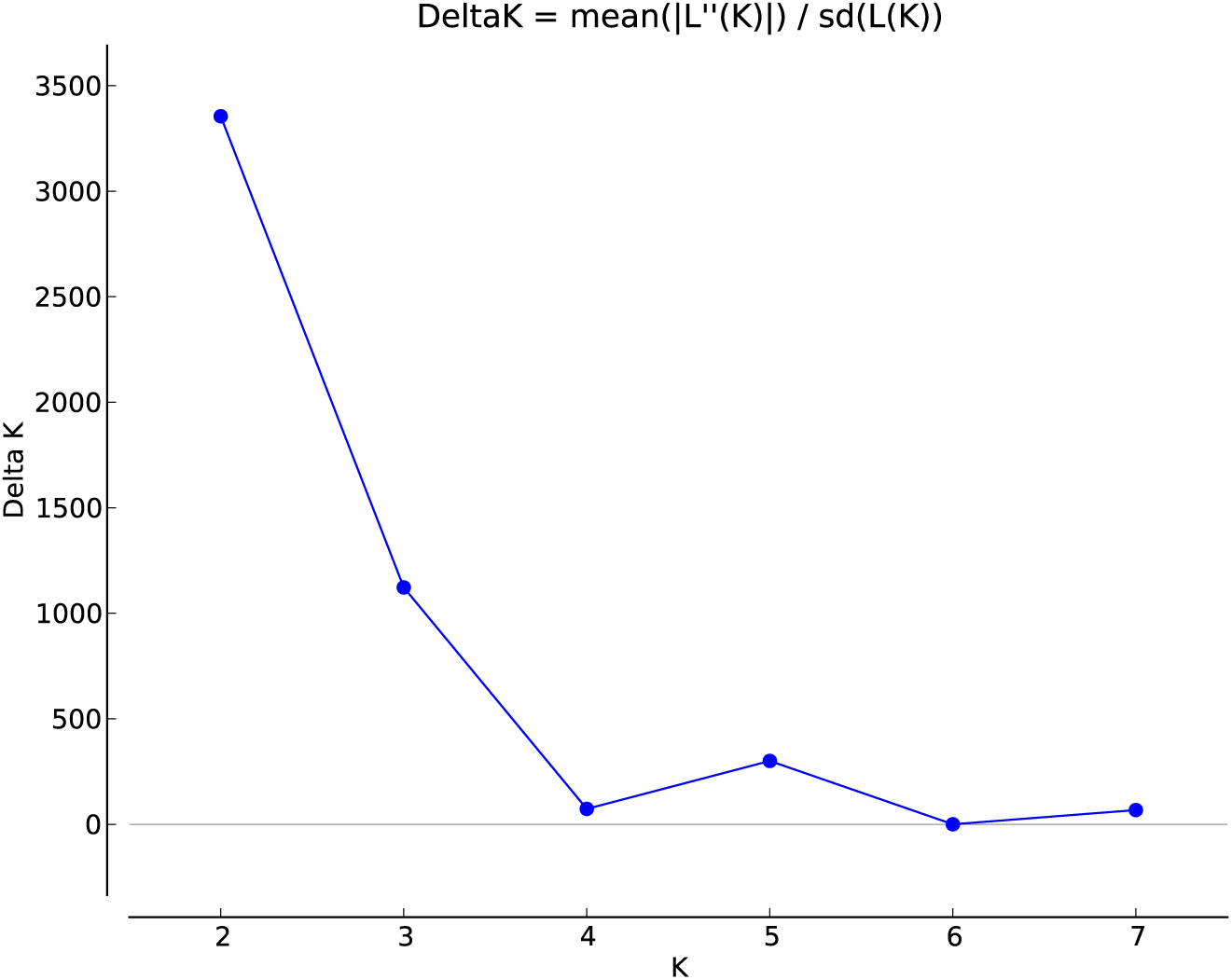
A plot of likelihood value for STRUCTURE analysis across a range of population numbers (*K*) from 2 to 7. The best fit to cowpea genotyping data occurs at *K*=5.

**Fig. S5.**
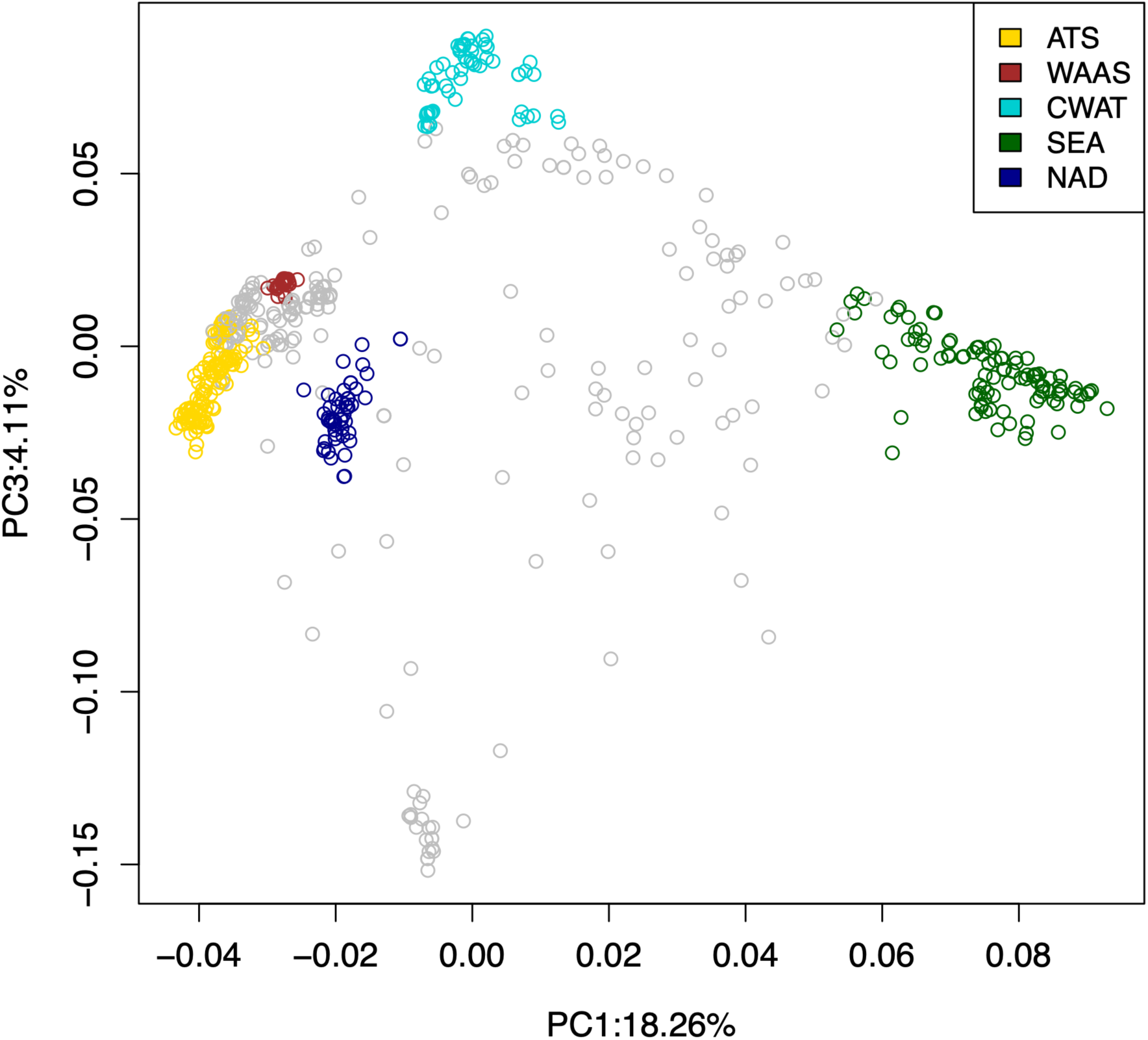
Principal Component Analysis (PCA) showing genetic clustering following Principal components 1 and 3. ATS = African Tropical Savannah (Yellow), WAAS = West African Arid Steppe (Red), CWAT = Coastal West African Tropical (Light blue), SEA = Southeastern Africa, (Green), NAD = North African Desert (Dark blue).

**Fig. S6.**
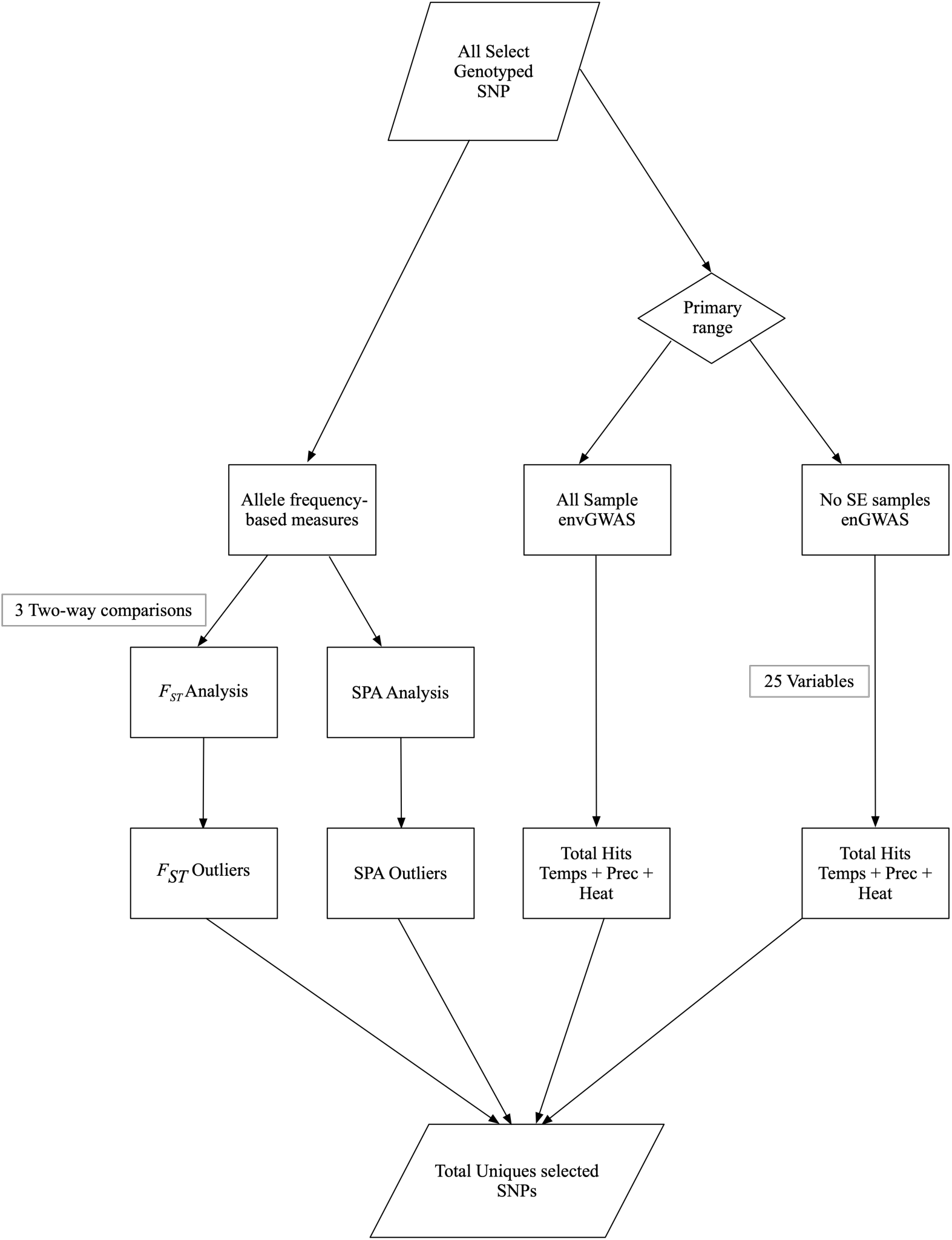
A workflow depicting outlier detection approaches used for detection of variants putatively associated with environmental adaptation in cowpea. The approaches in allele frequency comparisons (*F_ST_*), spatial ancestry analysis (SPA), and genome-wide association analysis using environmental variables (envGWAS).

**Fig. S7.**
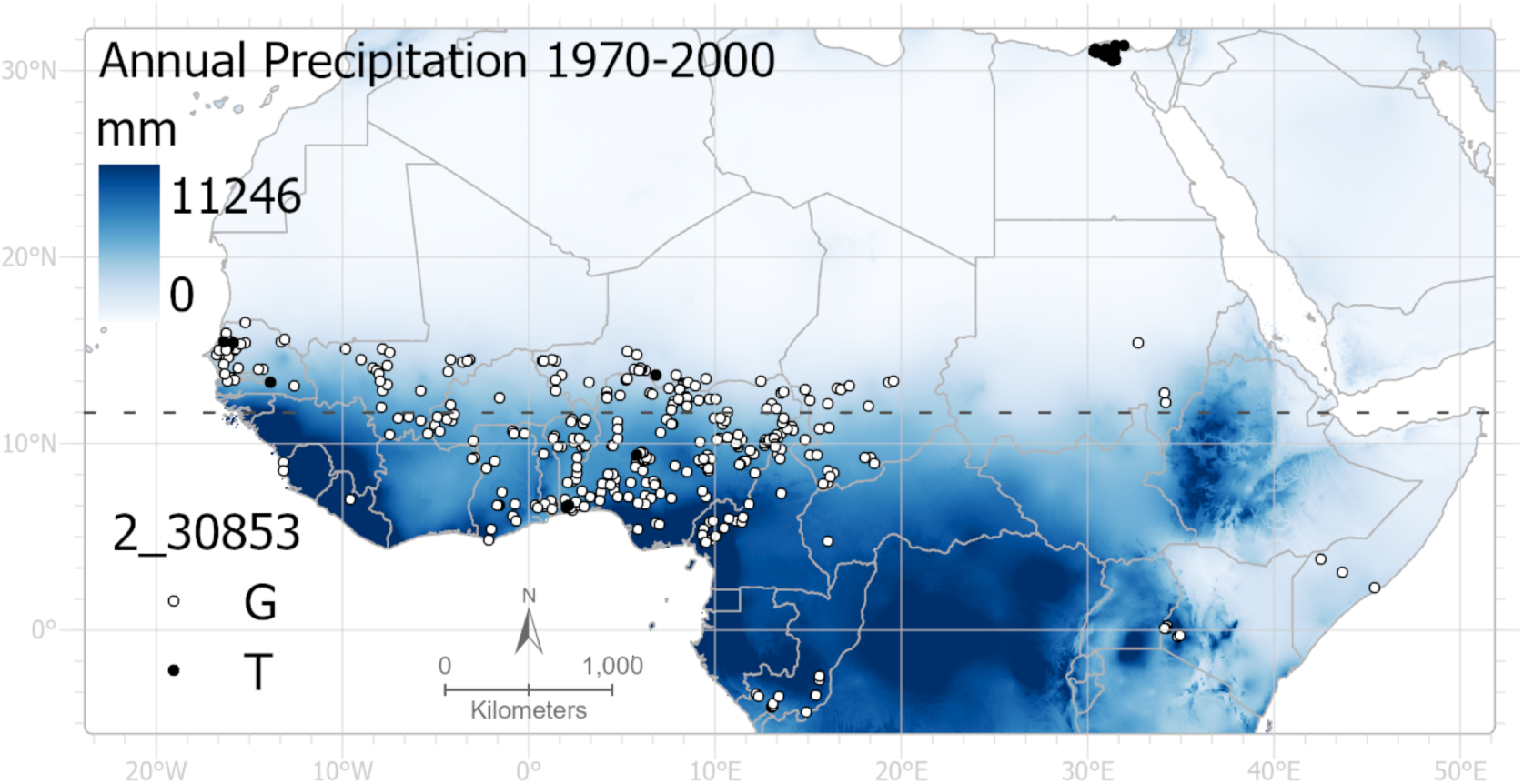
A comparison of the allele frequencies for the SPA outlier 2_30853 with annual precipitation. The map represents the annual mean precipitation from 1970 to 2000, with white indicating low precipitation and blue indicating high precipitation (ranging from 0 to 11250mm). Alleles for variant 2_30853 are plotted on the map: white dots represent the reference allele (G), and black dots represent the alternate allele (T).

**Fig. S8.**
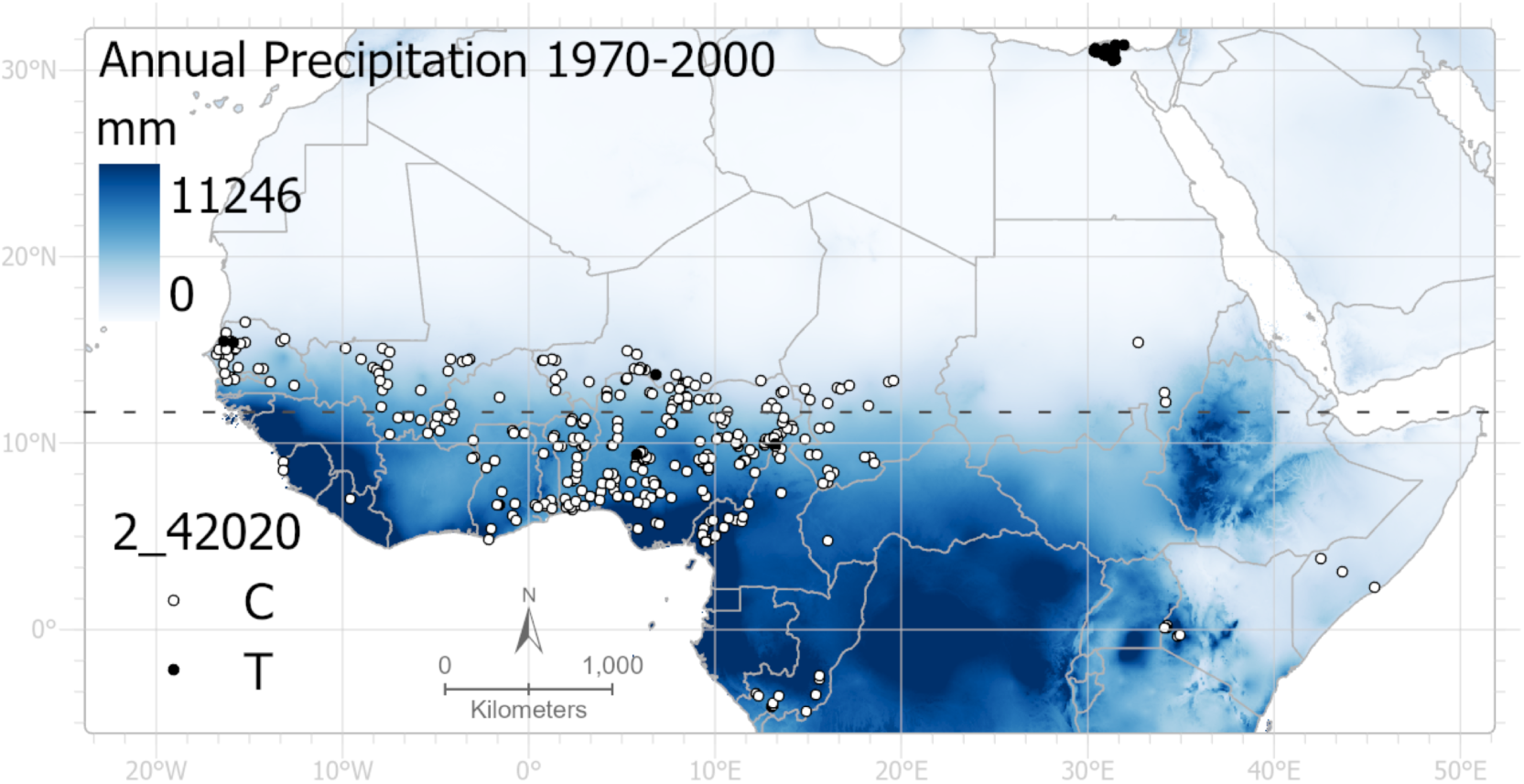
A comparison of the allele frequencies for the SPA outlier 2_42020 with annual precipitation. The map represents the annual mean precipitation from 1970 to 2000, with white indicating low precipitation and blue indicating high precipitation (ranging from 0 to 11250mm). Alleles for variant 2_42020 are plotted on the map: white dots represent the reference allele (C), and black dots represent the alternate allele (T).

